# Impact of diet on jejunal microbiota composition during broiler development with special focus on *Enterococcus hirae* and *Enterococcus faecium*

**DOI:** 10.1101/2023.04.05.532946

**Authors:** Paul B. Stege, Dirkjan Schokker, Frank Harders, Soumya K. Kar, Norbert Stockhofe, Vera Perricone, Johanna M. J. Rebel, Ingrid C. de Jong, Alex Bossers

## Abstract

Modern broiler breeds allow for high feed efficiency and rapid growth, but come at a cost of increased susceptibility to pathogens and disease. Broiler growth rate, feed efficiency, and health are furthermore affected by the composition of the gut microbiota, which in turn is influenced by diet composition. In this study we therefore assessed how diet composition alters the broiler jejunal gut microbiota. A total of 96 broiler chickens were divided into four diet groups: control, coated butyrate supplementation, medium chain fatty acid supplementation, or a high-fibre low-protein content. Diet groups were sub-divided into age groups (4, 12 and 33 days of age) resulting in groups of 8 broilers per diet per age. The jejunum content jejunum was used for metagenomic shotgun sequencing to determine the microbiota composition on species level. Among all diet groups, a total of 104 differential abundant bacterial species were detected. Most notably were the changes in the jejunal microbiota induced by butyrate supplementation when compared to the control diet, resulting in the reduced relative abundance of mainly *Enterococcus faecium* and the opportunistic pathogen *Enterococcus hirae* in broilers 4 days post-hatch. At this early stage of development, the immune system is still immature thereby highlighting the importance to study the relation of diet and the jejunal microbiota. Future studies should furthermore elucidate how diet can be used to promote a beneficial microbiota in the early stages of broiler development.

## Introduction

The continuous expansion of the poultry industry comes with the demand to improve sustainability of production. Modern broiler breeds offer the advantage of rapid growth and increased feed efficiency, but come with the disadvantage of increased susceptibility to physiological and metabolic disorders, and have indications of inferior immunity [1–6]. Although diet composition has also been optimized for sustainability in terms of growth rate and feed efficiency, the effect of diet on the composition of the intestinal microbiota has not been fully explored. Namely the composition of the jejunal microbiota and the potential diet-induced effects thereof are currently unknown, which is of specific interest as it is one of the principal sites of nutrient absorption [21].

The gastrointestinal tract and corresponding intestinal microbiota together contribute to feed efficiency, the development of the immune system and ultimately to the state of health and disease [7–10]. In turn, diet composition is known to alter both intestinal physiology and microbiota composition and is thus proposed as a tool to facilitate sustainability in terms of feed efficiency, animal health and reduced mortality. The beneficial effects of diet composition were first observed when growth promoters in the form of antibiotics were established to affect the intestinal microbiota, resulting in beneficial effects on the general health and feed efficiency of chickens [11–14]. However, this sub-therapeutic use of antibiotics has the added effect to enrich for antibiotic resistant bacteria and its use is now prohibited globally [15–20]. As a result, the impact of diet composition and additives on the intestinal microbiota are investigated in search for alternatives to mimic mostly the beneficial effects introduced by antibiotics. Although 16S rRNA gene sequencing remains the most common approach to determine diet-induced effects in the bacterial community composition, its resolution is surpassed by that of metagenomic shotgun sequencing (MSS). By sequencing the full microbiome, MSS is not only able to detect bacteria on the genera taxonomic-level but can additionally determine bacterial species. Moreover, MSS can be used to study gene composition and their corresponding gene pathways [21].

The small intestine is specialized for nutrient absorption, where amino acids are mainly absorbed in the proximal part of the jejunum, while fatty acids are utilized in the distal parts of the jejunum [22, 23]. Both parts are densely colonized with bacteria and in the case of broiler chickens, the most abundant bacteria in the small intestine include lactic acid-producing bacteria *Lactobacillus, Enterococcus* and *Streptococcus*, from which *Lactobacillus* is overall the most abundant genera [24–28]. The high abundance of *Lactobacillus* suggests that these bacteria play a prominent role in the intestines and is one of the reasons why *Lactobacillus* is commonly applied in probiotics [29, 30]. Diet composition is explored as an approach to induce shifts in the intestinal microbiota, for instance by altering the ratio of fatty acids and fibres in feed. When animal fat and soybean oil were supplemented with medium-chain fatty acids (MCFA; 0.3% C10 and 2.7% C12) for 34 days, the broiler ileum microbiota showed a reduction of *Lactobacillus*, *Enterococcaceae*, *Micrococcaeae* and an increase in *Enterobacteriaceae* [31]. MCFAs have been observed to have antibacterial properties against opportunistic pathogens like *Clostridium perfringens* and *Escherichia coli* when applied in *in vitro* experiments, but it is unknown if the antibacterial properties persist in a complex system as the intestinal microbiota [32–34]. Another example is butyrate, which is a short chain fatty acid (SCFA) and is the preferred energy-providing substrate of colonocytes [35]. When broiler feed was supplemented with butyrate for 42 days, both feed efficiency and villi size were increased [36]. Butyrate can be rapidly absorbed by the microbiota and intestinal cells located in the proximal sites of the intestine. In order to slowly release butyrate over the full length of the intestine, Mallo et al., 2021, supplemented coated butyrate for 42 days and observed similar results to uncoated butyrate [37]. Supplementation of fibre in feed is known to induce changes in the intestinal microbiota of broilers. Mainly the bacteria located in the caecal microbiota can ferment fibre, generating components including SCFAs [38]. While low level fibre supplementation can increase the amount of butyric acid in the cecum of 21-day-old broilers and increased the abundance of *Helicobacter pullorum* and *Megamonas hypermegale*, high levels of fibre supplementation increased the abundance of taxa that may include pathogens, namely *Selenomonadales*, *Enterobacteriales*, and *Campylobacterales* [39].

The majority of previously discussed studies analyse diet-induced effects on the genera taxonomic-level of bacteria, preventing the observation of for instance species-specific effects. Only a subset of bacteria has been used for species-specific comparisons, thereby potentially neglecting other important bacteria including opportunistic pathogens. Moreover, while the jejunum is one of the principle sites of nutrient absorption, the effects of diet on the jejunal microbiota are unknown. In this study, we therefore assessed how different diets impacts the composition of the jejunal microbiota on the species level by performing MSS of the microbiota in broilers during the first 33 days post-hatch.

## Results

### Diet-associated differences in the jejunal microbiota

A total of 96 Ross 308 broilers were housed in ground cages. They were divided into four diet groups, to study the effect of diet: (1) control diet (CON); (2) control diet supplemented with butyrate (BUT), (3) control diet supplemented with medium-chain fatty acids (MCFA) and (4) a diet with high-fibre low-protein composition (HFLP). The jejunal microbiota was studied by taking chyme samples after either 4, 12 or 33 days post-hatch, thus studying groups of 8 broilers per diet per timepoint. Samples were used for metagenomic shotgun sequencing, resulting in 137.6M [SE 67.4] reads per sample and 32.7M [SE 2.5] assigned read pairs per sample after taxonomic classification. Sample s2229 contained the lowest number of assigned read pairs (1.7M) and was therefore excluded from downstream analysis. This sample was part of the BUT group 12 days post-hatch. There was no systemic difference in the top 10 abundant bacterial species, between the diet groups. In all diet groups, the most abundant 10 species included: *Lactobacillus johnsonnii*, *Limosilactobacillus reuteri*, *Ligilactobacillus salivarius*, *Enterococcus hirae*, *Lactobacillus crispatus*, *Pediococcus acidilactici*, *Enterococcus faecium*, *Corynebacterium stationis*, *Limosilactobacillus vaginalis* and *Enterococcus faecalis*. These are all lactic acid bacteria, except for *C. stationis* [40, 41]. The jejunal microbiota displays an overall age dependent effect, independent of diet. This includes an overall high abundance of *E*. *hirae* 4 days post-hatch (relative abundance 12.95% [SE 0.04]), which is reduced at 12 days post-hatch (0.83% [SE 0.01]). In a similar way, the jejunal microbiota showed a steep decrease of *L*. *johnsonnii* from 12 days post-hatch (58.99% [SE 0.03]), compared to 33 days post-hatch (18.16% [SE 0.04]). In contrast, the abundance of *L. salivarius* and *L. crispatus* increased over the course of 12 days post-hatch (0.00% [SE 0.00]; 0.31% [SE 0.00]) to 33 days post-hatch (20.05% [SE 0.28]; 6.86% [SE 0.04]).

The jejunal microbiota diversity expressed as Shannon index and the microbiota evenness expressed as Pielou index, were not significantly different between diet groups (figure 2a, figure s1). Overall, the total species diversity is highly similar among diet groups. Multidimensional scaling (MDS) was applied on Bray-Curtis distance matrices and revealed that diet was not a main driver of the observed variance in microbiota composition between samples at either 4, 12 or 33 days post-hatch (figure 2b).

**Figure 1.**
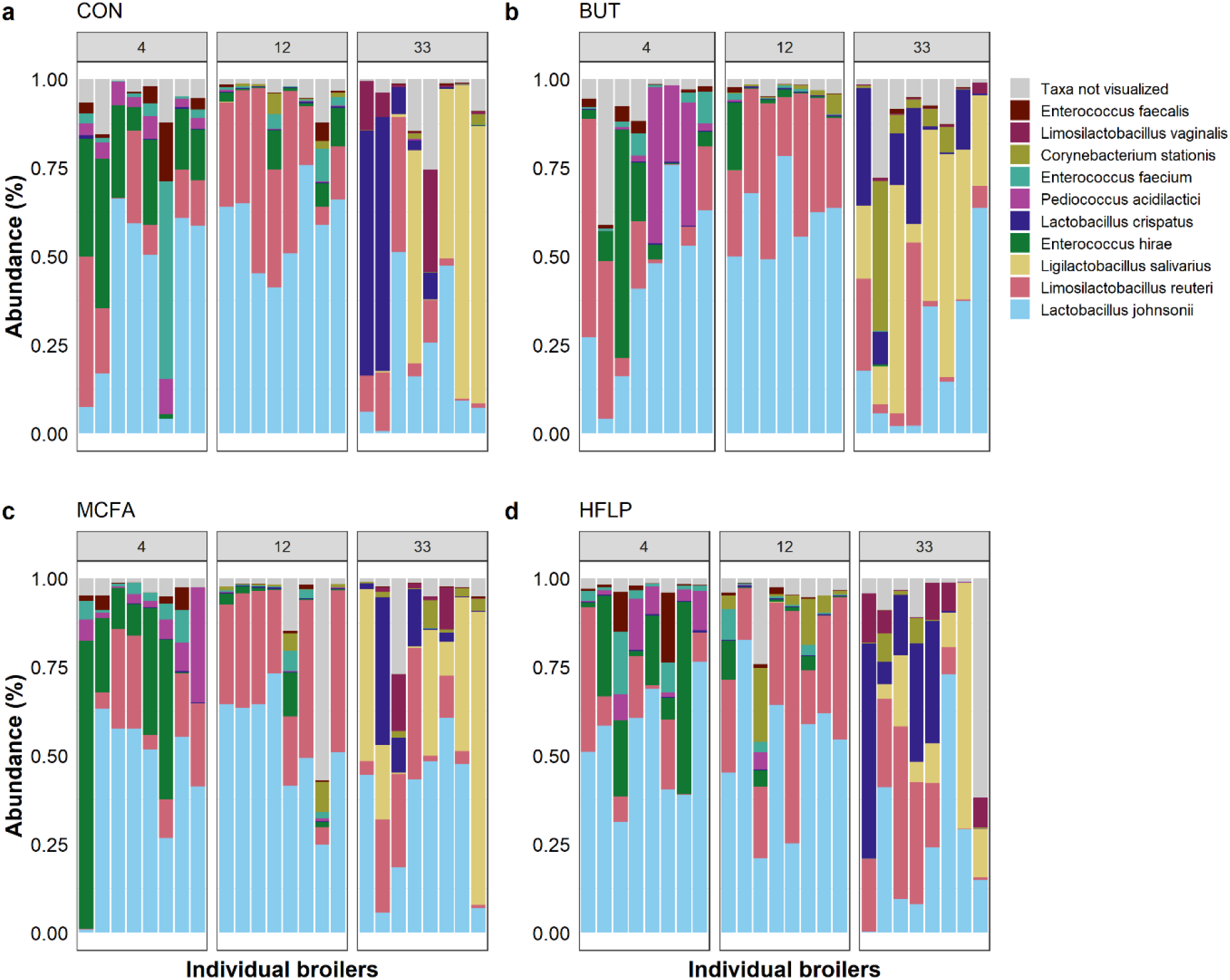
Association of diet and the composition of the jejunal microbiota at different ages (d4, d12, and d33). Relative abundance of the 10 most abundant bacterial species per diet group. (**a**) Control diet (CON), (**b**) control diet plus butyrate (BUT), (**c**) control diet plus medium-chain fatty acids (MCFA) and (**d**) a high-fibre low-protein diet (HFLP). Broilers are grouped by columns, representing the number of days post-hatch. Abundance was plotted on the relative abundance scale from 0.00 to 1.00.

**Figure 2.**
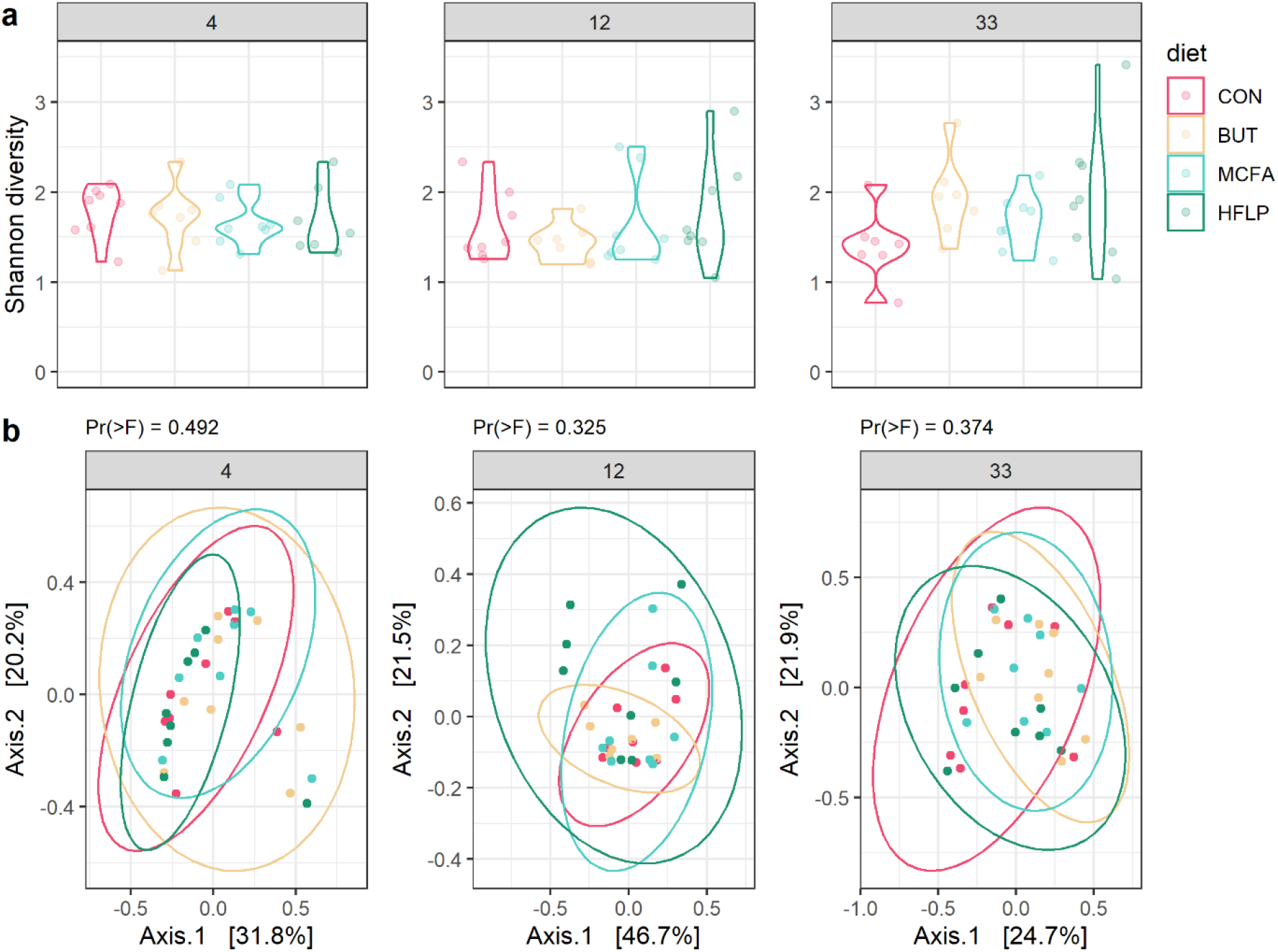
Diversity indices of jejunal microbiota per diet group at different ages (d4, d12, and d33). (**a**) Alpha diversity per diet group expressed by Shannon diversity index on OTU level. Diet groups did not differ in terms of alpha diversity when compared with Wilcoxon rank-sum tests. (**b**) Beta diversity of bacterial species, using MDS ordination based on Bray-Curtis dissimilarity on OTU level. Individual broilers and corresponding ellipses are coloured according to diet group. Plot panels represent broiler groups of 4, 12 and 33 days post-hatch. Permutational multivariate analysis of variance and testing for homogeneity of multivariate dispersions revealed no significant differences between groups.

Differential abundance analysis revealed a total of 104 bacterial species that were significantly different in terms of abundance when comparing diet groups to the control group (figure 4, table s1-9). At 4 days post-hatch, the jejunal microbiota of the **BUT** diet resulted in 43 differentially abundant bacteria, compared to the control group. Bacteria with a relative abundance above 0.01% and that changed in terms of relative abundance compared to the control group, expressed as log2 fold changes (l2fc), included: a reduction of *E. hirae* (−2.9 l2fc, 4.2% abundance), *Enterococcus faecium* (−1.8 l2fc, 1.2% abundance), *Enterococcus durans* (−2.6 l2fc, 0.04% abundance), *Erysipelatoclostridium ramosum* (−2.4 l2fc, 0.03% abundance), *Enterococcus avium* (−1.8 l2fc, 0.02% abundance), *Lacrimispora saccharolytica* (−1.5 l2fc, 0.01% abundance), *Massilistercora timonensis* (−1.5 l2fc, 0.01% abundance), *Lachnoclostridium phocaeense* (−1.4 l2fc, 8.3e-3% abundance) and of *Weissella paramesenteroides* (−5.1 l2fc, 2.2e-4% abundance). The **MCFA** diet resulted in 6 differentially abundant bacteria that are present in low abundance, including a reduction of *Pediococcus pentosaceus* (−1.6 l2fc, 0.04% abundance) and of *W*. *paramesenteroides* (−4.8 l2fc, 2.8e-4% abundance). The **HFLP** diet resulted in 7 differentially abundant bacteria that are present in low abundance, including a reduction of *P*. *pentosaceus* (−1.3 l2fc, 0.043% abundance) and of *W*. *paramesenteroides* (−5.1 l2fc, 7.3e-5% abundance).

**Figure 3.**
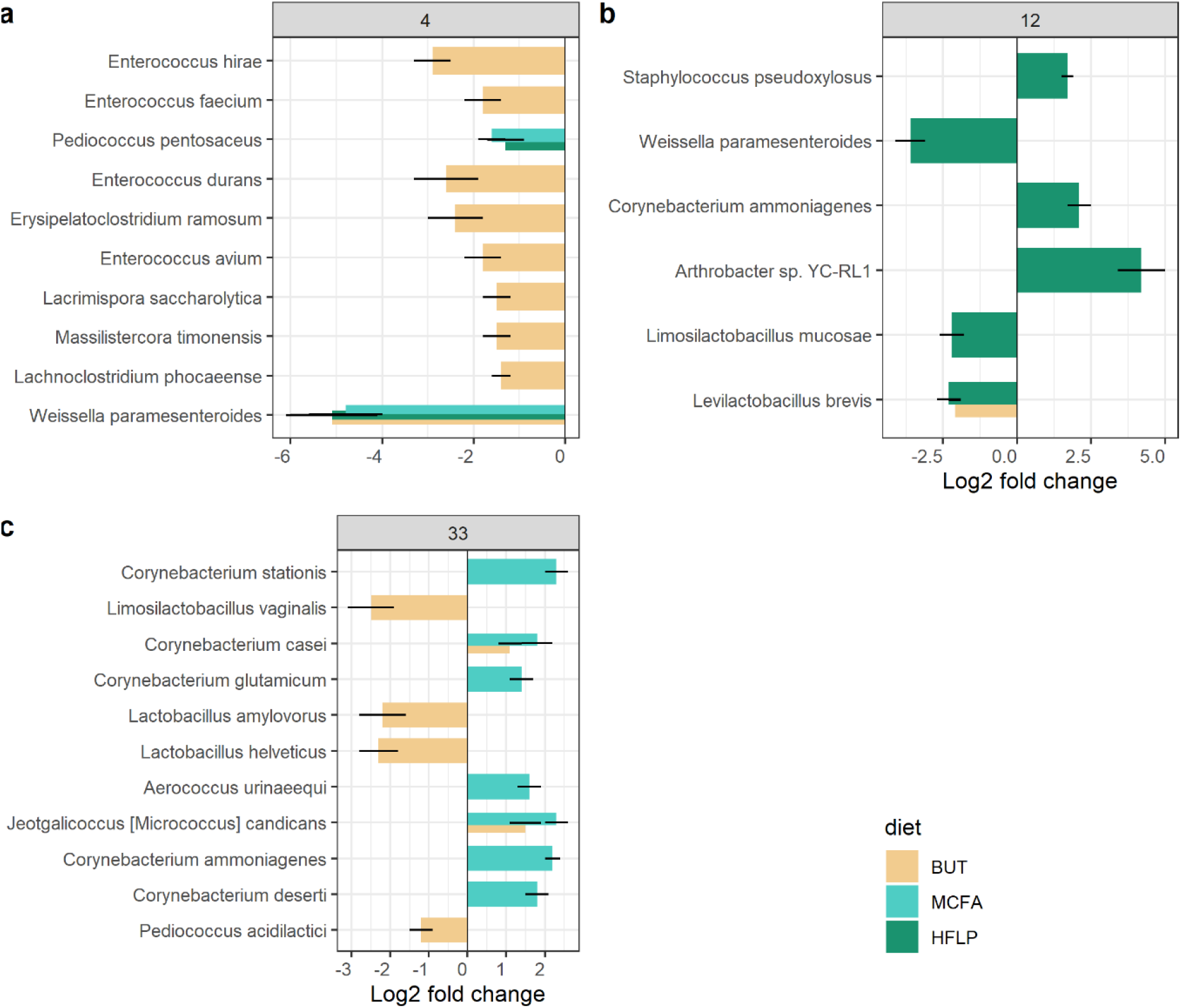
Diet-induced differentially abundant bacteria in jejunal microbiota at different ages (d4, d12, and d33). Differential abundance analysis on BUT, MCFA and HFLP diet groups as a function of the control diet group at (**a**) 4 days post-hatch, (**b**) 12 days post-hatch and (**c**) 33 days post-hatch. Log2 fold change (l2fc) differences are visualized by bars, coloured according to diet group and the standard error by error bars. Differential abundant bacteria are visualized (abundance > 0.001%; p-value < 0.05; l2fc > |1|) and ordered from most abundant (top) to least abundant (bottom).

**Figure 4.**
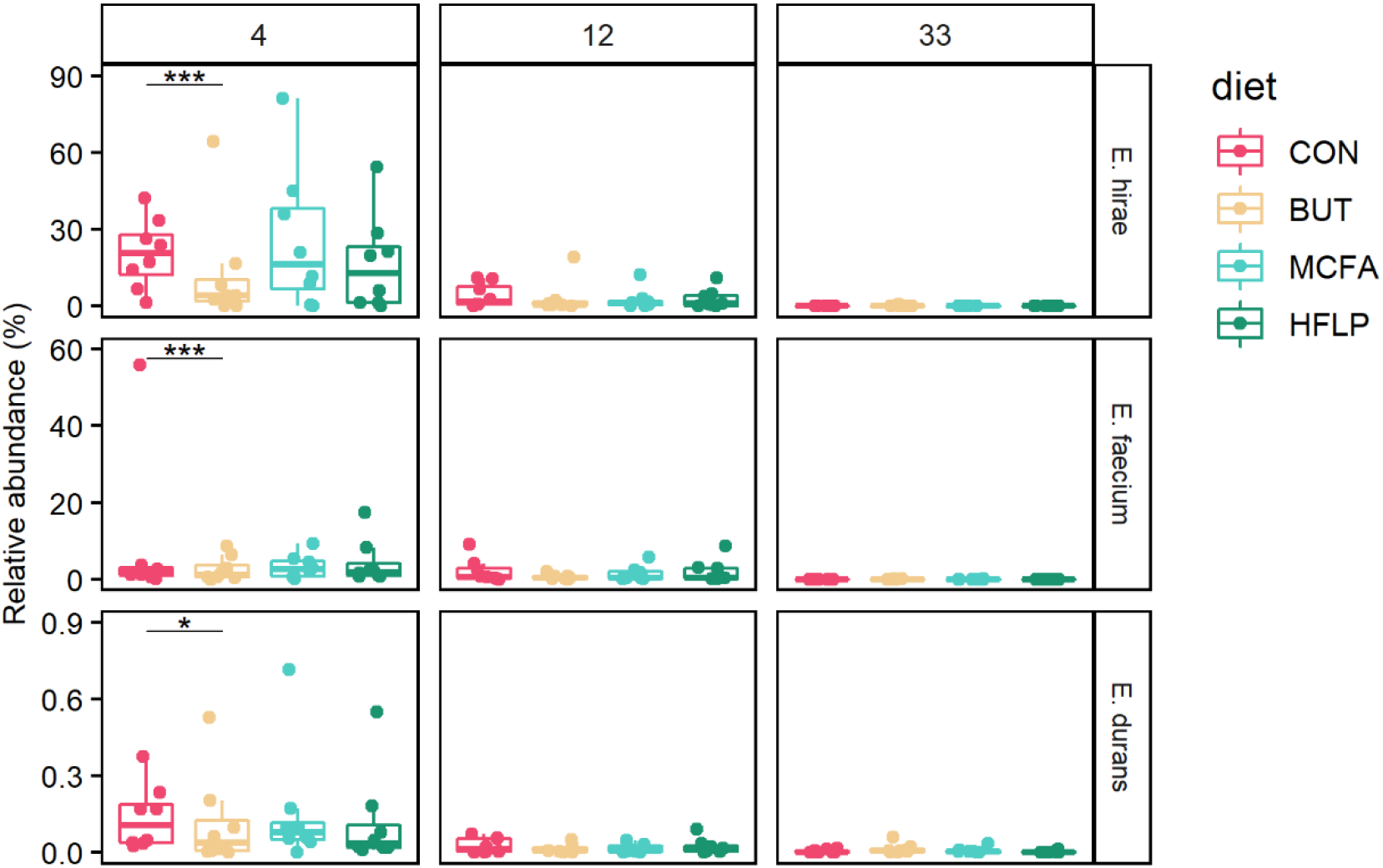
Diet-induced differentially abundant opportunistic pathogens. Relative abundance of opportunistic pathogens *E*. *hirae*, *E*. *faecium* and *E. durans*, indicated by rows. Columns represent broiler groups of 4, 12 and 33 days post-hatch. Individual broilers are coloured according to diet group. Adjusted p-values are calculated as part of ANCOMBC as function of the control diet and indicated by *< 0.05, **< 0.01 and ***< 0.001.

At 12 days post-hatch, the **BUT** diet resulted in 17 differentially abundant bacteria that are present in low abundance, including a reduction of *L. brevis* (−2.1 l2fc, 1.1e-3% abundance). The **MCFA** diet resulted in 3 differentially abundant bacteria that have a low relative abundance in the MCFA and control diet groups (all below 1e-2% abundance, table s5). The **HFLP** diet resulted in 31 differentially abundant bacteria, including the increase of *Staphylococcus pseudoxylosus* (1.7 l2fc, 0.71% abundance), *Corynebacterium ammoniagenes* (2.1 l2fc, 0.03% abundance), *Arthrobacter* sp. YC-RL1 (4.2 l2fc, 0.018% abundance and the decrease of *W. paramesenteroides* (−3.6 l2fc, 4.2e-4% abundance), *Limosilactobacillus mucosae* (−2.2 l2fc, 9.7e-5% abundance) and *L. brevis* (−2.3 l2fc, 1.4e-3% abundance).

At 33 days post-hatch, the **BUT** diet resulted in 18 differentially abundant bacteria, including a reduction of *L. vaginalis* (−2.5 l2fc, 0.42% abundance), *Lactobacillus amylovorus* (−2.2 l2fc, 0.07% abundance, *Lactobacillus helveticus* (−2.3 l2fc, 0.05% abundance), *P. acidilactici* (−1.2 l2fc, 0.01% abundance) and the increase of *Corynebacterium casei* (1.1 l2fc, 0.26% abundance) and *Jeotgalicoccus* (*Micrococcus*) *candicans* (1.5 l2fc, 0.03% abundance). The **MCFA** diet resulted in 33 differentially abundant bacteria, including the increased abundance of *Corynebacterium stationis* (2.3 l2fc, 1.7% abundance), *C*. *casei* (1.8 l2fc, 0.16% abundance), *Corynebacterium glutamicum* (1.3 l2fc, 0.12% abundance), *Aerococcus urinaeequi* (1.6 l2fc, 0.04% abundance), *J.* (*M.*) *candicans* (2.3 l2fc, 0.02% abundance), *C*. *ammoniagenes* (2.1 l2fc, 0.02% abundance) and *Corynebacterium deserti* (1.8 l2fc, 0.01% abundance). The **HFLP** resulted in 23 differentially abundant bacteria that have a low relative abundance in the HFLP and control diet groups (all below 1e-2% abundance, table s9).

### Diet-associated differences in the abundance of opportunistic pathogenic and potential beneficial bacteria

The observed diet-induced differentially abundant bacteria include the opportunistic pathogens *E*. *hirae*, *E*. *faecium*, *E*. *durans* and *S. pseudoxylosus*. From this selection, *E*. *hirae*, *E*. *faecium* and *E*. *durans* are present in high relative abundance in broilers in the control group at an early stage of broiler development (20.53% [SE 0.14], 1.86% [SE 0.19], 0.11% [SE 0.00] at 4 days post-hatch) compared to broilers of 12 days post-hatch (1.75% [SE 0.05], 0.66% [SE 0.03], 0.01% [SE 0.00], figure 4). *E*. *faecium*, *E*. *durans* and *E*. *hirae* were all shown to decrease in abundance at 4 days post-hatch as a result of butyrate supplementation compared to the control diet. In order to confirm that presence of these closely related species and exclude the possibility of misannotation by the aligner tool, sequencing data of the control diet group 4 days post-hatch was mapped to the genome of all detected enterococcal species (105.1M [SE 33.0] reads per sample). This resulted in a high number of reads per sample mapping to *E*. *hirae* (10.0M [SE 6.0] reads, 90.9% [SE 5.3] coverage, 415.7 [SE 286.1] depth), *E*. *faecium* (0.8M [SE 0.3] reads, 88.0% [SE 2.1] coverage, 31.4 [SE 15.8] depth), *E*. *faecalis* (0.3M [SE 0.3] reads, 88.6% [SE 6.6] coverage, 12.4 [SE 14.6] depth) and in lesser extend to other enterococcal species (all below 70% coverage), leading us to conclude that at least *E*. *hirae* and *E*. *faecium* are present in the jejunal microbiota of these broilers and affected by butyrate supplementation (table s10).

This analysis was repeated for the bacteria *L*. *mucosae*, *L*. *vaginalis*, *L*. *brevis*, *L*. *amylovarus, L*. *helveticus*, *P*. *pentosaceus* and *W*. *paramesenteroides* since these are considered beneficial to gut health and applied in probiotics [42–47]. *L*. *mucosae* and *L*. *brevis* were shown to decrease in abundance at 12 days post-hatch as a result of the HFLP diet compared to the control diet. At 12 days post-hatch butyrate supplementation results in a decreased abundance of *L*. *brevis*. Sequencing data of the control diet group 12 days post-hatch was mapped to the genome of all detected *Limosilactobacillus* and *Levilactobacillus* species (106.9M [SE 4.5] reads per sample). This resulted in a high number of reads per sample mapping to *L. reuteri* (7.3M [SE 1.6] reads, 82.1% [SE 0.4] coverage, 403.1 [SE 100.2] depth) while other detected *Limosilactobacillus* and *Levilactobacillus* species had a coverage below 70%. Due to the low number of reads and coverage, the differential abundance of *L*. *mucosae* and *L*. *brevis* at 12 days post-hatch could therefore not be confirmed. *L*. *vaginalis*, *L*. *amylovarus* and *L*. *helveticus* were shown to decrease in abundance at 33 days post-hatch as a result of the BUT diet compared to the control diet. Sequencing data of the control diet group 33 days post-hatch was mapped to the genome of all detected *Limosilactobacillus* and *Lactobacillus* species (113.9M [SE 35.0] reads per sample). This resulted in a high number of reads per sample mapping to *L. reuteri*, (1.8M [SE 2.0] reads, 83.6% [SE 5.9] coverage, 103.7 [SE 128.5] depth), *L. vaginalis*, (0.4M [SE 2.2] reads, 86.8% [SE 10.9] coverage, 26.2 [SE 140.5] depth) and to *L*. *crispatus* (2.7M [SE 14.4] reads, 87.1% [SE 8.3] coverage, 154.1 [SE 934.3] depth), while other detected *Limosilactobacillus* and *Lactobacillus* species had a coverage below 70%. Due to the low number of reads and coverage, the differential abundance of *L*. *vaginalis*, *L*. *amylovarus* and *L*. *helveticus* at 33 days post-hatch could therefore not be confirmed.

*P*. *pentosaceus* was shown to decrease in abundance at 4 days post-hatch as a result of MCFA and HFLP diet. *P*. *acidilactici* on the other hand, decreased in abundance at 33 days post hatch as a result of the BUT diet. When mapping the sequencing data of the control diet group at 4 and 33 days post-hatch to all detected *Pediococcus* species (105.1M [SE 33.0] and 106.9M [SE 4.5] reads per sample, 4 and 33 days post-hatch respectively). This resulted in a high number of reads per sample mapping to *P*. *acidilactici* 4 days post-hatch (1.3M [SE 1.4] reads, 87.2% [SE 1.0] coverage, 76.3 [SE 92.5] depth), while other detected *Pediococcus* species at 4 or 33 days post-hatch had a coverage below 70%. Due to the low number of reads and coverage, the differential abundance of *P*. *pentosaceus* at 4 days post-hatch and *P*. *acidilactici* 33 days post-hatch could therefore not be confirmed.

Finally, *W*. *paramesenteroides* was shown to decrease in abundance at 4 days post hatch for the BUT, MCFA and HFLP diet and at 12 days post hatch for the HFLP diet. When mapping the sequencing data of the control diet group at 4 and 12 days post-hatch to all detected *Weisella* species (105.1M [SE 33.0] and 106.9M [SE 4.5] reads per sample, 4 and 12 days post-hatch respectively). The detected *Weisella* species at 4 or 12 days post-hatch had a coverage below 70% and therefore the differential abundance of *W*. *paramesenteroides* at these timepoints cannot be confirmed.

## Discussion

In this study, we determined the jejunal bacterial microbiota of broilers 4, 12 and 33 days post-hatch using metagenomic shotgun sequencing (MSS) and evaluated to what extend diet modulates the jejunal microbiota composition. The results confirmed that diet supplemented with either butyrate (BUT), medium chain fatty acids (MCFA) or diet with high-fibre low-protein (HFLP), can induce significant differences in the relative abundance of a total of 104 bacterial species. The results of butyrate supplementation are of specific interest as they reduced the relative abundance of highly abundant enterococci in the jejunal microbiota 4 days post-hatch, a critical stage for broiler health. Specifically, MSS allowed to differentiate between bacteria on species level and revealed that butyrate supplementation greatly reduces the relative abundance of *Enterococcus hirae* and *Enterococcus faecium*.

Regardless of the fluctuations in microbiota composition in the first weeks of life, we observed that the most abundant species are mainly part of the genera *Lactobacillus*. This is in concordance with previous studies that analysed the small intestines of broilers [24, 27, 48]. MSS allowed us to surpass the level resolution of 16S rRNA gene sequencing. To our knowledge, this is the first time that the composition of broiler jejunal microbiota has been determined at the bacterial taxonomic species level. When comparing the ten most abundant bacterial species at 4, 12 and 33 days post-hatch, we observed fluctuations in the jejunal microbiota during broiler development. This included a transition from high relative abundant *Lactobacillus johnsonii* (58.99% [SE 0.15]) at 12 days post-hatch to *Ligilactobacillus salivarius* and *Lactobacillus crispatus* 20.35% [SE 0.28]; 6.86% [SE 0.21]) at 33 days post-hatch. This is similar to the findings of Lu et al., 2023, when studying the broiler ileum microbiota. They observed a transition of the most dominant species, switching from *Lactobacillus acidophilus* at 14 days post-hatch (53% abundance) to *L. crispatus* at 28 days post-hatch (75% abundance) [24]. The ileum is the small intestinal region located directly downstream of the jejunum and the corresponding microbiotas are known to share similarities in their composition, potentially explaining these similar findings [49].

We observed *E. hirae* to be the second most abundant bacterial species in the jejunal microbiota 4 days post-hatch among diet groups (12.95% [SE 0.21]) and observed a much lower relative abundance of *E*. *hirae* 12 days post-hatch (0.83% [SE 0.05]). This matches with the findings of Schokker et al., 2017, where a decrease in overall enterococcal abundance was observed from 21.7% 4 days post-hatch to 4.9% 14 days post-hatch [48]. We determined that supplementation of butyrate to broiler feed resulted in a 2.9 log2 fold change decrease in abundance of *E*. *hirae* in broilers 4 days post-hatch. Additionally, the butyrate supplemented group showed a decrease of several enterococci species at 4 days post-hatch, including *E. faecium* and *Enterococcus durans* and *Enterococcus avium*. From these, only *E*. *hirae* and *E*. *faecium* were present in sufficient abundance to ensure that this species is present with at least 70% genome coverage. The butyrate applied in this study is coated, which was previously found to result in the slow release of butyrate along the length of the intestinal tract [50, 51]. Sun et al., 2022, has shown that coated butyrate can reduce the abundance of enterococci in the ileum microbiota of squabs [52], but this is the first time this function is demonstrated in broilers. *E*. *hirae* is an opportunistic pathogen that can cause locomotion problems, endocarditis and septicaemia in broiler chickens [53–56]. The effect of butyrate on the abundance of *E*. *hirae* 4 days post-hatch is of specific interest, since this early stage represents the most critical period during broiler development. In this stage, the immune and digestive system are still immature, thereby increasing the susceptibility to disease [57]. This period is furthermore marked by the transition from aerial breathing, initiation of thermal regulation and changes in diet composition, from yolk to solid feed, contributing to the overall high stress load during early broiler development [58]. While some short chain fatty acids directly inhibit bacterial growth, butyrate supplementation only resulted in limited growth inhibition of *E*. *faecium* and *E*. *hirae*, when tested *in vitro* [32–34, 59]. Therefore, this effect is not likely to be caused by butyrate directly, but rather indirect, i.e. by changes in the jejunal microbiota as a result of the butyrate supplementation.

*E*. *faecium* is known as a gut commensal, but also as an opportunistic pathogen in humans and animals [60, 61]. Moreover, specific isolates of *E*. *faecium* are applied as probiotics In broilers [62, 63]. The distinction between these groups lies in differences in their genetic makeup [64, 65]. The detected *E*. *faecium* should therefore first be classified in order to conclude about its impact on broiler health. *L*. *mucosae*, *L*. *vaginalis*, *L*. *brevis*, *L*. *amylovarus*, *L*. *helveticus*, *P*. *pentosaceus* and *W*. *paramesenteroides* are considered beneficial for gut health and were found differentially abundant as a result of diet. Due to the low relative abundance, these species did not reach the threshold of 70% genome coverage and their presence could therefore not be validated.

Modern broilers breeds have been primarily selected for rapid growth rate and feed efficiency. While diet composition has additionally been optimized for broiler growth rate, feed efficiency and health benefits, the relation between diet and the jejunal microbiota is not fully understood. In this study we showed that supplementation of diet with either BUT, MCFA or HFLP induced changes in the jejunal microbiota composition at bacterial species level of broilers 4, 12 or 33 days post-hatch. Most notably was the observed reduction in the abundance of *E*. *faecium* and of the opportunistic pathogen *E*. *hirae* in the BUT diet, compared to the control diet. The impact of diet composition on microbiota composition of opportunistic pathogens during early broiler development, highlights the importance to study the relation of diet and the jejunal microbiota. Future studies should furthermore elucidate how diet can be used to promote a beneficial microbiota in the early stages of broiler development and could be supported by metatranscriptomics.

## Methods

The experiment was conducted at the experimental research facility of Wageningen University and Research. All procedures complied with the Dutch law on animal experiments; the project was approved by the Central Commission on Animal Experiments (license number AVD4010020197985) and the experiment by the Ethical Committee of Wageningen University & Research, the Netherlands; experiment no. 2019.D-0009.001.

### Classification of broiler groups

Day-old Ross 308 male broiler chickens were obtained from a commercial hatchery (Probroed & Sloot, Groenlo, The Netherlands) and housed floor pens with wood shavings as substrate *ad libitum* access to feed and water as described by Perricone et al., 2023 (REF PENDING). Broilers were divided into age groups of 4, 12, and 33 days post-hatch, which were subdivided into four diet groups for microbiota analysis, as described in Perricone et al., 2023 (REF PENDING). In summary, these groups are: (1) control diet without any supplementation (CON); (2) control diet supplemented with sodium butyrate (BUT, Excential Butycoat®, Orffa, Werkendam, the Netherlands), (3) control diet supplemented with a mixture of medium-chain fatty acids (MCFA, Aromabiotic®, Nuscience, Belgium) and (4) a diet with a high-fibre low-protein composition compared to the control diet (HFLP). While the BUT and MCFA diet involve supplementation of components, the HFLP diet involved substitutions of several components of the control diet, including the substitution of rapeseed meal by potato protein and an increase of sunflower seed meal and corn, and a reduction in soybean meal. To summarize, 8 broilers were studied per diet per timepoint. Cages were separated into 8 blocks (1 broiler/diet group/age/block), in order to prevent the exchange of manure and or litter, as described by Perricone et al., 2023 (REF PENDING). Feed was provided via a round feeder (diameter: 35 cm) hanging in the pen. Water was provided via seven nipples along the side wall of a pen. Broilers were vaccinated against infectious bronchitis before arrival at the experimental facility and on d25, and against Newcastle disease at d15.

### Sample collection, storage and DNA extraction

Chyme samples were collected from the jejunum of broilers by squeezing the jejunum content in a collection tube. Samples were snap-frozen and transferred to storage at −80°C. One freeze-thaw cycle was introduced when dividing samples into aliquots of 0.2 g. Aliquoted samples were used for DNA extraction with the Invitrogen PureLink Genomic DNA Mini Kit according to the manufacturer’s instructions. Total DNA was quantified by using an Agilent 2200 Tapestation.

### Metagenomic shotgun sequencing and data processing

DNA samples were sent to GenomeScan B.V. (Leiden, the Netherlands) for Metagenomic shotgun sequencing. The NovaSeq 6000 (Illumina, San Diego, USA) was applied with S2 flow cells and the 2 x 150 bp paired-end kit (Illumina) according to company protocols. Samples contained on average 137.6M [SE 67.4] reads. Sequencing reads were adapter-clipped, erroneous-tile filtered, and quality-trimmed at ≥ Q20 (PHRED score) using BBduk v38.96 and subsequently filtered for host DNA using the global-alignment algorithm of BBmap v38.96 with default settings and *fast=t* (broiler genome version 2021/01/19, accession number GCF_016699485.2) [66]. Read pairs were then used for taxonomic classification by Kraken v2.1.2 using the premade standard Kraken RefSeq nucleotide database and applying a confidence cut-off of 0.3 (database version 5/17/2021) [67]. This resulted into 32.7M [SE 2.5] assigned read pairs per sample. The sample with the lowest number of assigned read pairs (1.7M) was excluded from downstream analysis (s2229, BUT group 12 days post-hatch). Kraken2 results were converted into a biom-file using kraken-biom v1.0.1 at standard settings in order to export counts on strain level, here referred to as OTU level [68].

### Data analysis

Analysis of sequencing data was performed in R version 4.0, RStudio v2022.02.2+485 and functions of R packages phyloseq (version 1.4) and ggplot2 [69, 70]. The top 10 abundant bacteria in the jejunal microbiota were plotted using the tax_glom function of the phyloseq package (while removing unassigned reads), aggregate function of microbiomeutilities and plotting functions of the microbiome package [71, 72]. The Shannon diversity and Pielou evenness index were calculated on OTU level by first applying rarefaction to an equal library size (720,000 reads, matching the sample with the lowest number of reads), using the rarefy_even_depth function of phyloseq (set.seed=194175, replace=FALSE) [69]. Consequently, the alpha diversity function of the microbiome package was applied [72]. The beta diversity analysis using MDS ordination and Bray-Curtis dissimilarity was calculated on OTU level using the distance and ordinate functions of the phyloseq package [69]. Permutational Multivariate Analysis of Variance (PERMANOVA) and tests on homogeneity of dispersion were employed using the adonis2 function (999 permutations, seed of 194175) and betadisper function of the vegan package [73]. Differential abundance analysis was performed by first selecting bacterial species for 10% prevalence and 0.001% abundance and subsequent analysis using ANCOM-BC version 1.6.0. ANCOM-BC was applied with standard settings, including Bonferroni correction for false discovery rate, batch correction for cage blocks and an alpha of 0.05 [74]. Structural zeros were included in the analysis (*struc_zero=TRUE*) and are indicated in tables s1-s9. A subsequent cut-off of 0.01% abundance per bacterial species per diet group and an absolute fold change cut-off of 2 were applied to generate the differential abundance plot (figure 3). The validation of detected bacterial species was performed by listing the reference genomes from all enterococcal species detected by kraken and downloading the corresponding RefSeq sequence from the NCBI database, filtering on full genomes and selecting the top hit when sorting by significance. Potential plasmids were excluded from the reference genomes and the resulting genomes were used to create a database using KMA version 1.4.3 and the index function with settings *-sparse TG*. Sequencing reads were subsequently aligned to this database using KMA and settings *-1t1*, *-ca*, *-apm p* and *-ef* [75].

### Data availability

Sequencing files have been submitted into the Sequence Read Archive (SRA) at the NCBI under accession number PRJNA952340. The phyloseq object is available at 10.5281/zenodo.7744071.

### Funding

This research was funded by the WUR program 2019-2022 KB34 Towards a Circular and Climate Neutral Society.

## Supporting information

Table_s1-9 - Differentially abundant species

Table_s10 - control_diet_species_mapping

## Supplementary information

**Figure s1.**
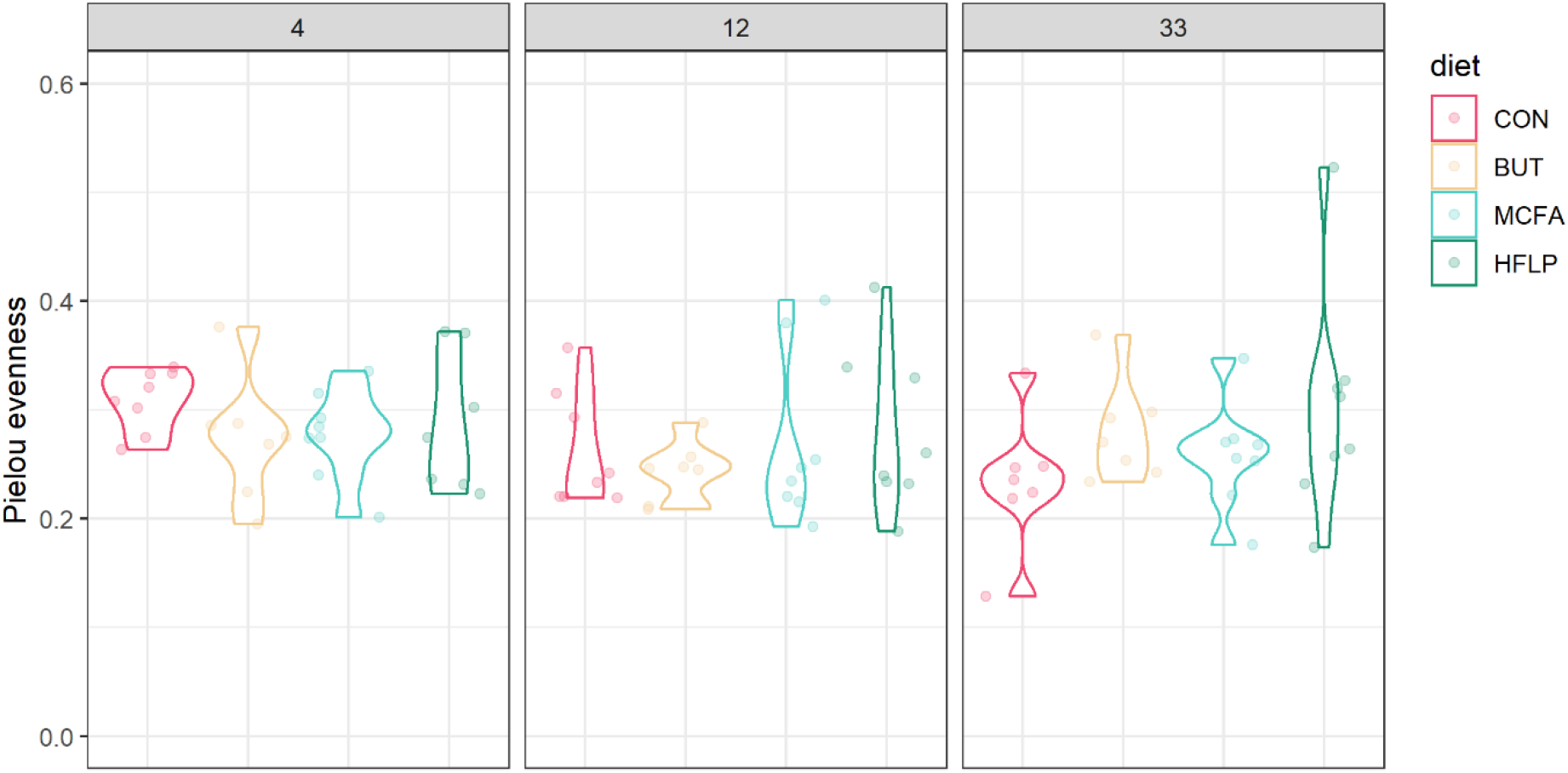
Microbiota evenness per diet group. Alpha diversity per diet group expressed by Pielou evenness index on OTU level. Diet groups did not differ in terms of alpha diversity when compared with Wilcoxon rank-sum tests.

## Notes

### Competing Interest Statement

The authors have declared no competing interest.

